# Single-Cell Analysis of Hyperthermic Intraperitoneal Chemotherapy Treated Tumors Reveals Distinct Cellular and Molecular Responses

**DOI:** 10.1101/2020.10.05.326710

**Authors:** Max P. Horowitz, Zahraa Alali, Tyler Alban, Changjin Hong, Emily L. Esakov, Tae Hyun Hwang, Justin D. Lathia, Chad M. Michener, Robert DeBernardo, Ofer Reizes

## Abstract

Hyperthermic intraperitoneal chemotherapy (HIPEC) has emerged as a clinical regimen that prolongs overall survival for patients with advanced Epithelial Ovarian Cancer (EOC). However, the mechanism of action of HIPEC remains poorly understood. To provide insights into the rapid changes that accompany HIPEC, tumors from five patients with high grade serous ovarian cancer were harvested from the omentum at time of debulking and after 90 minutes of HIPEC treatment. Specimens were rapidly dissociated into single cells and processed for single cell RNA-seq. Unbiased clustering identified 19 cell clusters that were annotated based on cellular transcriptome signatures to identify the epithelial, stromal, T and B immune cells, macrophages, and natural killer cell populations. Hallmark pathway analysis revealed heat shock, metabolic reprogramming, inflammatory, and EMT pathway enrichment in distinct cell populations upon HIPEC treatment. Collectively, our findings provide the foundation for mechanistic studies focused on how HIPEC orchestrates the ovarian cancer tissue response.

## Introduction

Epithelial ovarian cancer (EOC) is the 5^th^ most common gynecologic cancer with an average survival of two years. In 2018 approximately 22,000 women were diagnosed in the United States and the majority of them will ultimately succumb to their disease (1). The reason underlying the poor survival is due to poor diagnosis with 80% of patients with EOC present in advanced stage (III-IV) accounting for the poor prognosis (5-year cancer-specific survival 42% and 26%, respectively) (1). The mainstay of therapy is “optimal surgery” leaving minimal residual disease in combination with platinum and taxane based chemotherapy. While most women with advanced disease enter remission, recurrence is common and the majority of patients ultimately develop resistance to platinum-based therapy and succumb to their disease (2). There are few therapies in recent years that have been shown to significantly improve OS for patients with advanced or recurrent EOC. In a subset of EOC patients with recurrent platinum-sensitive disease who have tumors harboring BRCA1/2 mutation or homologous recombination-deficient (HRD) abnormalities, Poly ADP Ribose Polymerase Inhibitors (PARPi) have been shown to prolong overall survival (3). However, the only form of chemotherapy that has consistently been shown to significantly improve OS in the primary setting has come from administration of platinum- and taxane-based therapy directly into the abdominal cavity (IP therapy) (4). Despite significant improvement in outcomes and National Comprehensive Cancer Network (NCCN) recommendations, adoption of IP therapy is complex, requires overnight hospitalization and is associated with increased toxicity thus limiting its widespread adoption (2). Hyperthermic intraperitoneal chemotherapy (HIPEC) overcomes many of the issues with conventional IP therapy. HIPEC is chemotherapy that is heated to 42^°^C and administered into the abdomen once the surgical resection of tumor is completed. In a well-designed randomized controlled trial, the addition of HIPEC at time of interval debulking surgery in newly diagnosed ovarian cancer patients was shown to extend OS by nearly 12 months as compared to patients receiving identical treatment without HIPEC (5). A second randomized controlled trial demonstrated significant improvement in both PFS and OS in women with EOC that undergo surgery and HIPEC at recurrence with patients receiving HIPEC surviving 26.7 months compared to 13.4 months with surgery alone. Interestingly, patients that have developed resistance to platinum agents when given intravenously have been shown to respond to HIPEC using a platinum drug with survival outcomes equal to those with platinum responsive tumors (6).

Despite its proven clinical benefit in newly diagnosed patients as well as patients with both platinum-sensitive and -resistant cancers, the mechanism of action of HIPEC is unclear. Is the improvement in tumor control driven by increased direct cytotoxic effects or alterations in the tumor microenvironment? What role does the immune system play? Much of what is thought to be driving this effect is speculative and remains poorly understood. Understanding how HIPEC is improving outcomes is critical to further optimize treatment for patients with advanced EOC and will potentially shed light on targetable pathways that are currently not appreciated.

Mouse and rat studies have focused on establishing models to elucidate the impact of hyperthermia on EOC (7, 8). These have yielded limited insights on mechanisms and potential therapies. In parallel, cell based studies focused on applying hyperthermia on cultured cells (9) were able to identify altered molecular pathways that provide insights into the efficacy of HIPEC in humans. However, these data are limited given they are performed in an environment in isolation without a full impact of the tumor microenvironment and the impact of a functional immune system. To overcome this limitation, we analyzed ovarian tumors at the time of the debulking surgery and immediately following HIPEC protocol. As EOC is often found to metastasize to the omental fat, we focused on this site to interrogate cellular and molecular changes. We leveraged single cell RNA sequencing to identify the major cellular populations within the omental tumors (the tumor microenvironment) and underlying cellular and molecular changes accompanying HIPEC. Our data demonstrate that HIPEC activates immune cells and modulates the transcriptome of epithelial and stromal cell populations in patients with advanced EOC.

## Results

### Single Cell RNA Sequencing and Clustering Reveals Heterogeneity of EOC in Omentum

We utilized omental matched pre- and post-HIPEC tumor tissue from five patients for single cell RNA sequencing (Figure 1). All patients were diagnosed with high-grade serous ovarian cancer and underwent surgery at the Cleveland Clinic (Table 1). To date, 2 patients remain in remission, 2 recurred, and 1 is deceased. Standard quality control exclusion was applied, and a total of 69,200 cells from pre and post HIPEC tissues were analyzed. Unbiased cellular clustering resulted in identification of 19 unique clusters, which were consistent among the ten specimens, as visualized based on graph-based Uniform Manifold Approximation and Projection (UMAP) (Figure 2A). As the paired tissues were collected at different dates due to the patients’ surgeries, we validated that post-HIPEC specimen were representative of pre-HIPEC specimen using overlay UMAP to test for similarity in cluster distribution of the paired samples (Figure 2B).

**Table 1.** Patient demographics, tumor characteristics, and current status.

| Patient | A | B | C | D | E |
| --- | --- | --- | --- | --- | --- |
| Age at Surgery | 64 | 86 | 61 | 63 | 68 |
| Tumor Type | High Grade Serous Carcinoma | High Grade Serous Carcinoma | High Grade Serous Carcinoma | High Grade Serous Carcinoma | High Grade Serous Carcinoma |
| Stage | IIIC | IIIB | IIIC | IIIC | IIIC1 |
| Grade of Tumor | 3 | 3 | 3 | 3 | 3 |
| Tumor Location | Omentum | Omentum | Omentum | Omentum | Omentum |
| Ethnicity | Not-Hispanic | Not-Hispanic | Not-Hispanic | Not-Hispanic | Not-Hispanic |
| Race | White | White | White | White | African American |
| Co-morbidities at Surgery | Depression; Hypertension | Breast Cancer (1982); Cardiac (Pacemaker-A-fib); Hypertension; | none | Hypertension; Hypothyroidism | Stroke; Hypertension; Depression |
| status | Remission | Recurrence-pelvic and CA125 rising | Recurrence-scans and CA125 rising | Remission | Recurrence-scans and CA125 rising |
| Alive/Dead | Alive | Alive | Alive | Alive | Dead |

**Figure 1.**
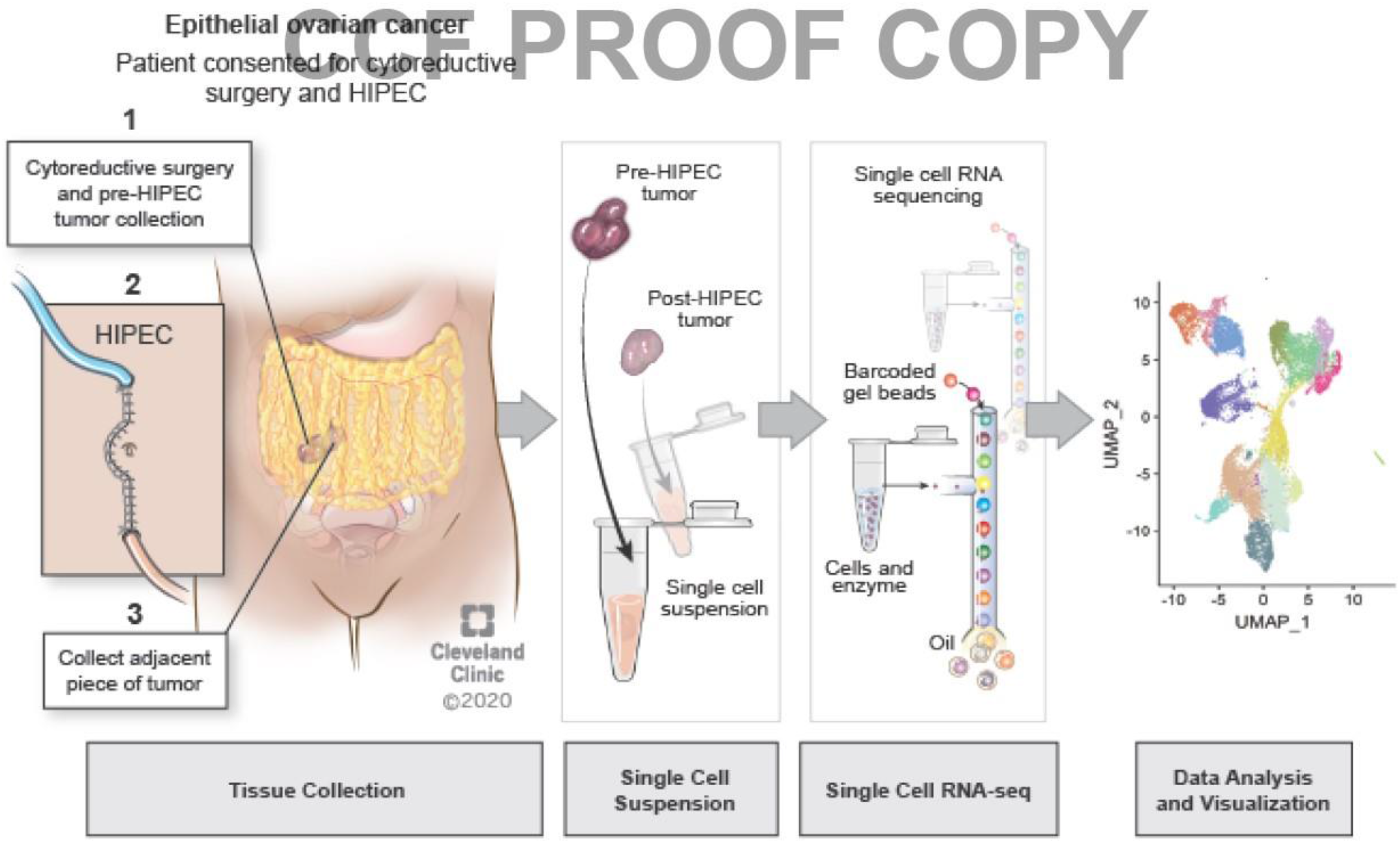
Schematic representation of the flow of tissues collection and processing for single cell RNA sequencing. A tumor specimen is obtained during debulking surgery and immediately post 90 minute HIPEC administration. Specimens are obtained from tumor in the omentum in order to reduce variability. Tumor samples (pre HIPEC and post HIPEC) are dissociated individually into a single cell suspension, and then the samples were loaded onto chromium column to generate barcoded cDNA. scRNA-seq 3’ libraries are generated and data are analyzed using The Cell Ranger. The final data are visualized into UMAP clustering and heatmap.

**Figure 2.**
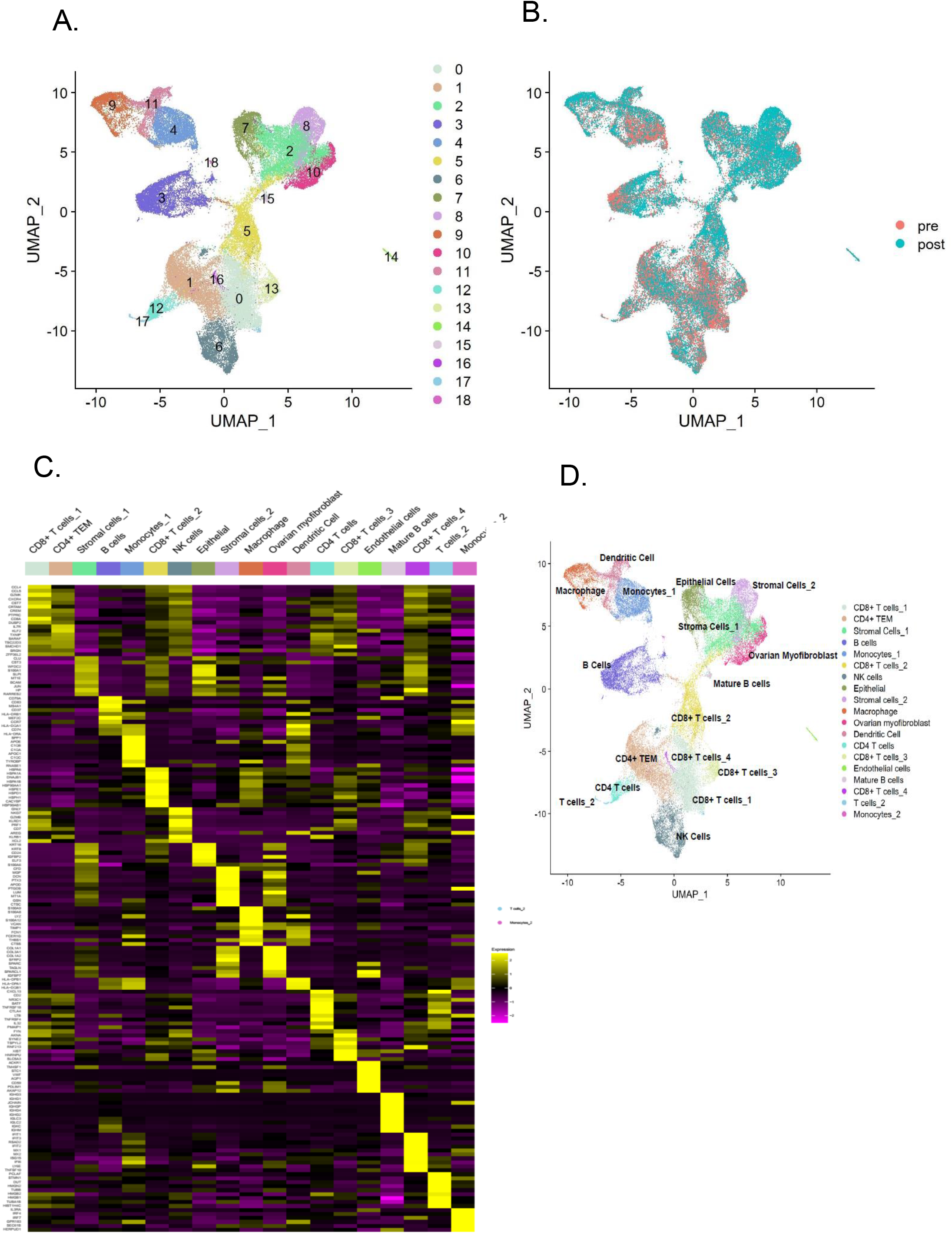
Single cell RNA sequencing analysis reveals tumor heterogeneity. (A) UMAP plot of tumor specimen combined from all pre and post HIPEC tissues. Unbiased clustering separates cells into 18 clusters based on gene expression profile. n=10 Samples Clustered from n=5 patients pre and post treatment Identify 19 distinct cell populations. (B) Overlay UMAP plot with integration of tumor tissues before and after treatment. n=10 Samples colored by pre-treatment and post-treatment samples. (C) Heatmap of canonical cell markers used for cell type annotations in tumor samples, color coded for gene expression level (yellow to black). (D) UMAP projection of cell populations showing the distribution of cell types and sub-types identified with each dot representing individual cell.

We next annotated the cell populations that comprised the unique clusters based on the top differential gene expression of established canonical cell markers (see Methods section) (Figure 2C, Supplemental Figure 1, Supplemental Table 2). Single cell RNA sequencing analysis detected heterogeneous cell populations distributed in each cluster (Figure 2D). Myeloid cells were identified in clusters 4, 9, and 11; B-cells in clusters 3 and 15; T cells in clusters 5, 1, 16, 12, 0, 7, and 13; and epithelial and stromal cells are found in clusters 7, 8, and 2. We then measured the percentage of each cell types in matched samples and found that immune cells are highly represented in the pre- and post-HIPEC specimens (Figure 3A, Supplemental Figure 2).

**Figure 3.**
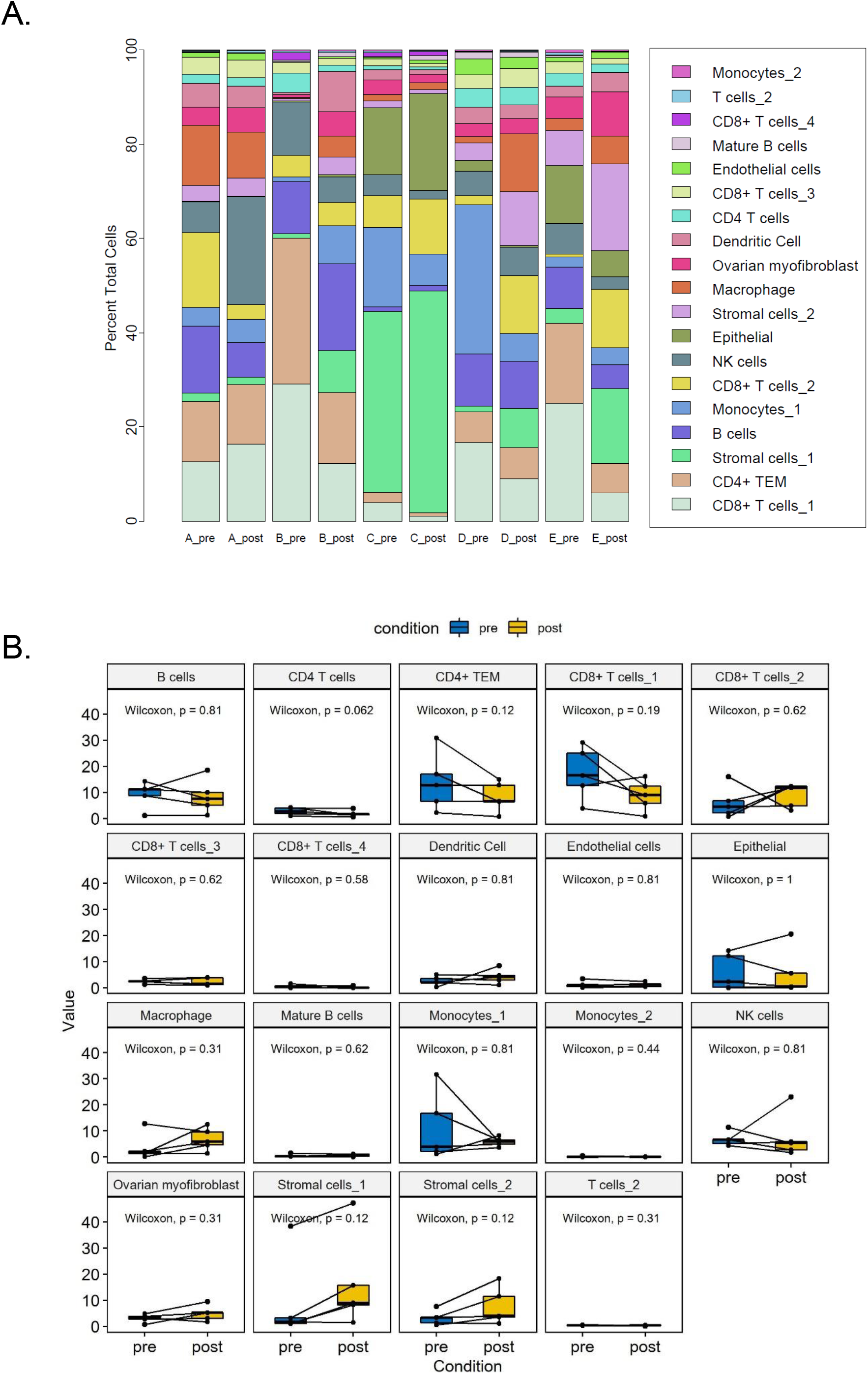
Relative abundance of infiltrating cells in tumor samples in response to HIPEC. (A) Bar plot representing the “percentage” of cell populations identified in each sample in tissues collected before and after HIPEC treatment of patients A, B, C, E, and D. (B) Plot depicting fraction of cell populations in post HIPEC tissues normalized to the pre-HIPEC samples per cell type.

We observed that the proportion of each individual cell type between pre-treated and matched HIPEC samples did not significantly change among cells types (Figure 3B; t test <0.05). Therefore, we focused on how HIPEC impacts cell populations at the transcriptional level within the immune, myeloid, epithelial, and stromal cells, comparing pre-treatment to post-treatment, with the hypothesis that HIPEC induces rapid transcriptional changes that can be observed after 90 minutes of treatment.

### Bulk Analysis Identifies Activation of Epithelial Mesenchymal Transition (EMT) Pathways and Upregulation in Inflammation Associated Genes

To investigate the differential gene expression (DGE) in pre- and post-HIPEC omental tissues, we utilized computational Gene Set Enrichment Analysis (GSEA). We found treated tissues share a unique genomic signature that potentially resulted from the induction of heat and chemical stress. Ninety minutes of HIPEC treatment was sufficient to induce significant changes in the expression of thousands of genes (Supplemental Table 3) with the top upregulated and downregulated genes represented in a volcano plot, with cutoffs at, p < 10^−16^, and Log2 fold change of 100. (Figure 4A). Notably, a list of genes found involved in inflammation, tumor suppression, and hyperthermia sensitivity are over-expressed in treated tissues compared to pre-treated tumors including *PTX3* (10-12), *CTGF* (9, 13), and *IGFBP7* (14, 15) (Figure 4A). The GSEA Hallmark pathway gene set annotations for the differentially expressed genes show activation of fatty acid metabolism, hypoxia, and epithelial mesenchymal transition (EMT) post-treatment (Figure 4B). These findings complement studies pointing to the tumor microenvironment and immune cells in directing and controlling tumor progression and chemotherapy response (16).

**Figure 4.**
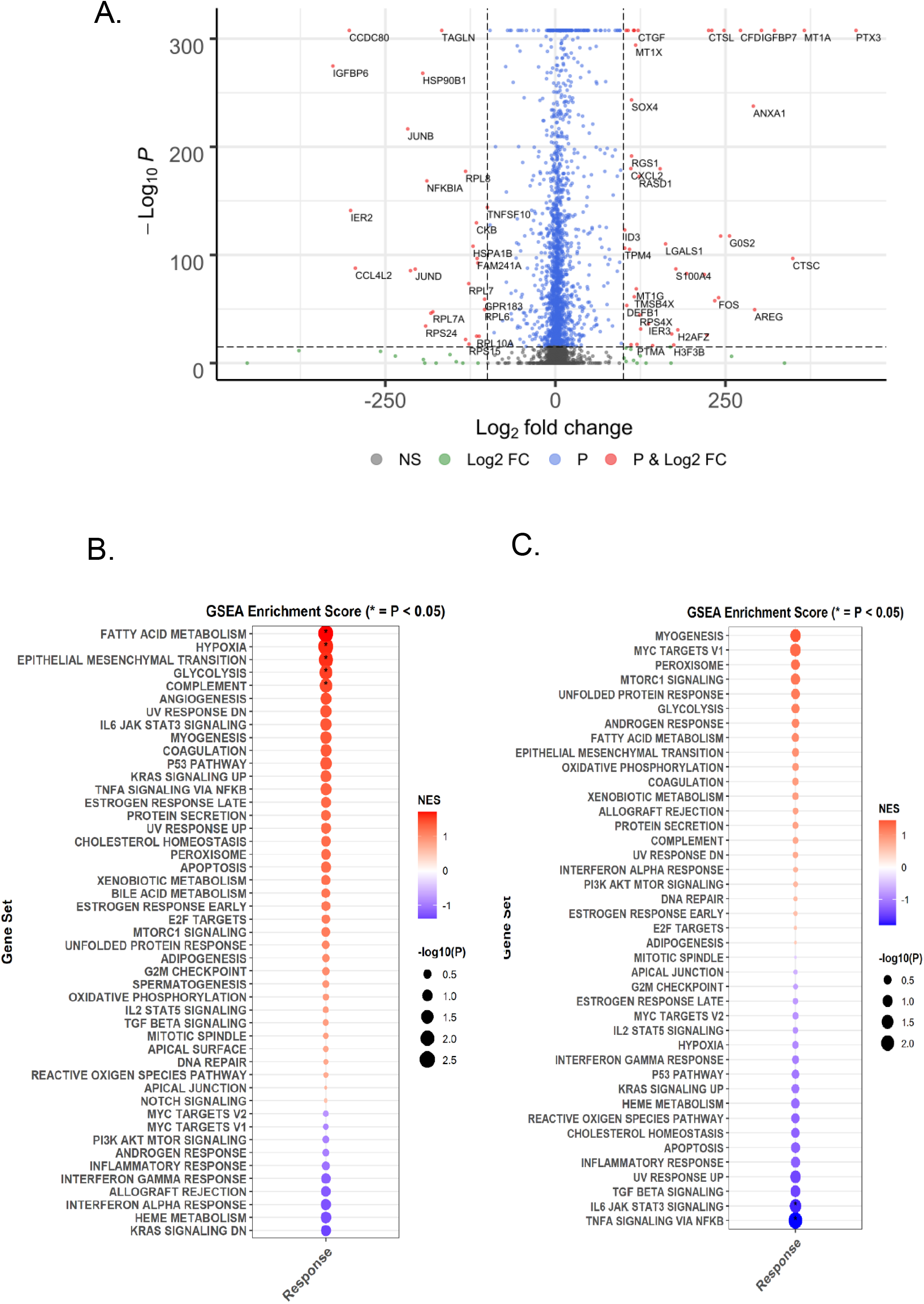
Genes enrichment and pathways analysis in treated tissues reveals induction of inflammatory and immune response. **(A)** Volcano plot of differential expressed genes in all cluster post HIPEC compared pre HIPEC tissues. Upregulated genes are represented on the right, and downregulated genes are shown on the left of the plot. (B) Dotplot depicting functional enrichment analysis of the positively activated (red) and negatively activated (blue) pathways in all clusters post HIPEC tissues. (C) Dotplot represent the pathways enriched in T cells collected after HIPEC. (from red to blue). Star denotes significant enrichment (p <.05).

**Figure 5.**
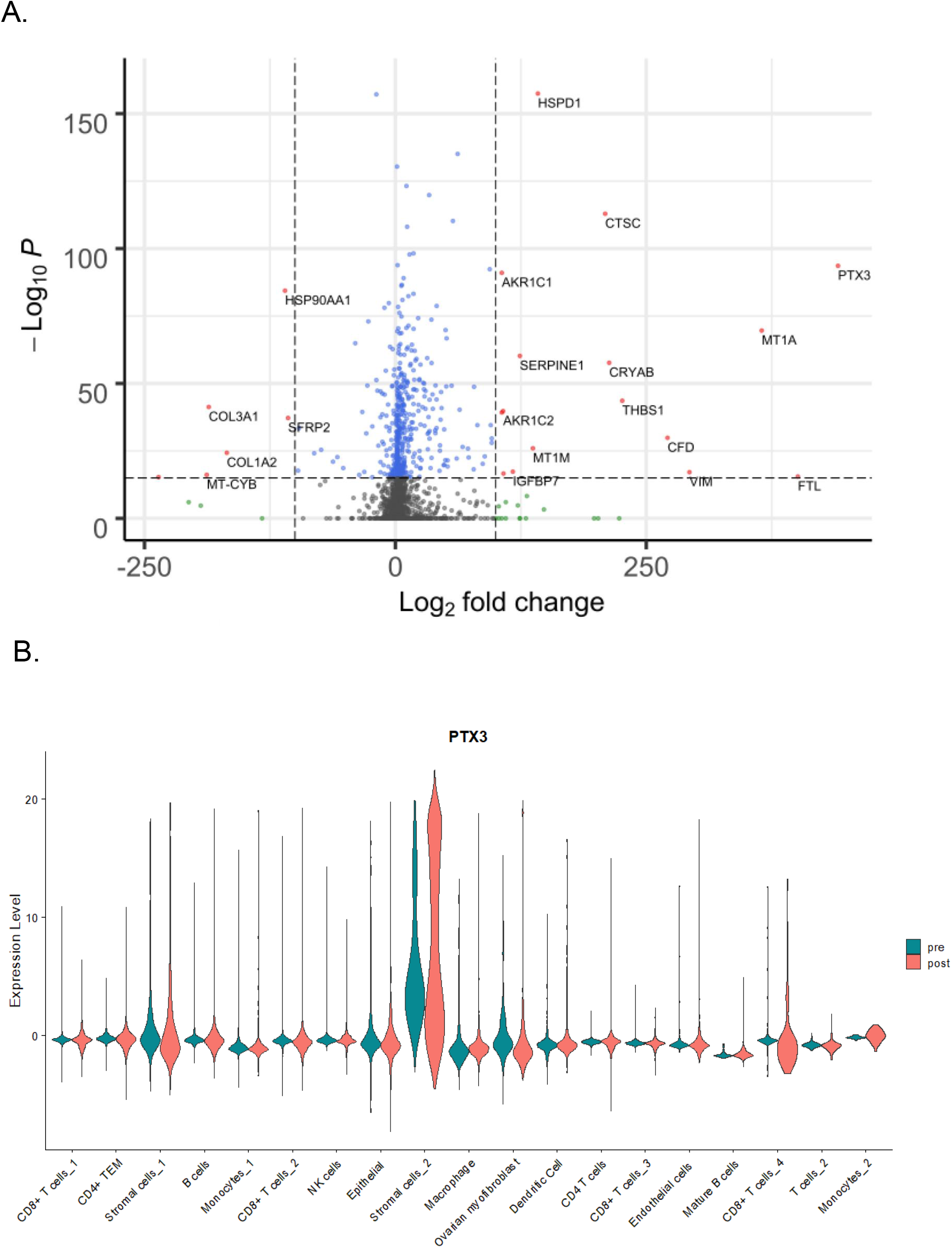
Molecular characterization and pathways analysis of stromal cells. (A) Volcano plot of the top and bottom differential expressed transcripts in post treatment tissues compared to pre HIPEC samples in stromal cells, represented by average fold change of expression and p-value. (B) Violin plot of the distribution of PTX3 expression in all identified population showing the abundance of PTX3 in stromal cells.

Furthermore, KRAS signaling and interferon alpha response are found to be inhibited in comparison to pre-treatment specimen. (Figure 4B). This analysis is equivalent to bulk RNAseq and cannot attribute the specific DGE to the complex landscape of the tumor microenvironment undergoing HIPEC. We then used the power of single cell RNA sequencing to identify the unique cell composition based on transcriptomic profiles followed by DGE analysis to identify genes and pathways altered by 90 minutes of HIPEC treatment.

### T- Lymphocytes and Natural Killer Cells Activate Global Cellular Differentiation and Propagation Pathways Post-HIPEC

We identified 7 sub-clusters of T cell populations, each labelled based on the expression of unique transcriptomic signature. CD8+ T cells are abundant in cluster 0, 5, 13, and 16 and were named CD8+ T cells_1, CD8+ T cells_2, CD8+ T cells_3, and CD8+ T cells_4 that differ in their gene expression profile (Figure 2A-D, Supplemental Table 2). CD4+ T cells are found in cluster 1 and 12. Cluster 1 includes the memory effector cells (CD4+ TEM), which express significantly high level of *IL7R* (interleukin-7 receptor), an enhancer for T cell survival and proliferation (17). Cluster 12 is a distinct sub-type of CD4+ that over-express *CXCL3* (Supplemental Table 2) known to attract B cells in severely inflamed tissues and aids in antibody productions (18). Natural killer cells were found in cluster 6 that express *CD69, KLRB1, GZMH, FCGR3A* (Figure 2A2D).

To identify the transcriptomic profile of treated T lymphocytes compared to the non-treated populations, we used differential gene expression analysis and identified genes involved in ribosomal biogenesis including *RPS8, RPS12, RPL41*, and *RPS2* (Supplemental Table 4). T lymphocyte activation involves cell cycle changes known to prepare cells for cellular differentiation and propagation. Using the differentially expressed genes of lymphocytes post treatment, we performed GSEA analysis that detected positive activation of myogenesis, as well as Myc target responsive genes (Figure 4C). We also identified significant negative activation of cytokine regulation pathways: TNF and IL6 signaling mechanisms (Figure 4C). Together, these data indicate HIPEC treatment leads to the transcriptional activation and expansion of lymphocytes early on during therapy.

### CD83 Tumor-Infiltrating-B Cells Exhibit Activation of Heat Shock and Estrogen Receptor Signaling in HIPEC Treated Tumors

Single cell analysis of B cells revealed the existence of a large population of B cells that overexpress the canonical cell markers *MS4A1, BANK1*, and *CD79A* in cluster 3 and a small sub-set of mature B cells (*IGKC+*) distributed in cluster 15 (Figure 2A-D, Supplemental Table 2).

DGE analysis of treated B-lymphocytes showed a transcriptomic signature that includes significant upregulation of heat shock proteins *HSPA5* and *HSPB1*, a consequence of hyperthermia treatment (19-24) as well as induction of mitochondrial transcripts *MT-CO1, MT-ND1*, and *MT-ND2* in post-HIPEC specimen compared to pre-treated specimen (Supplemental Table 5). The expression of mitochondrial genes is linked to B lymphocyte development, maintenance, and activation following stimulation of the adaptive immune response (25, 26). Notably, B-lymphocytes demonstrated a significant elevation in *CD83* expression (P <.05, fold change >1.5) (Supplemental Table 5), a marker of B cell activation, which is enhanced by antigen-specific T cell stimulation (27, 28).

GSEA pathway analysis was used to elucidate the physiological pathways activated in B lymphocytes cell clusters indicated activation of metabolic pathways and demonstrated a significant enrichment in estrogen response pathways (Supplemental Figure 3A). Here, we identified a significant increase in the *SPI-1* transcription factor and estrogen responsive gene that facilitates B cells response to external stimuli (25). Our data revealed that HIPEC stimulates the humoral immune response through triggering estrogen signaling and glycolysis pathway, which is known to result in reactivation of B-lymphocytes.

### HIPEC Stimulates MTORC1 Signaling Pathway in Myeloid Cell Population and Alters Tumor-Promoting Macrophage M2 Density

Cellular annotation of myeloid cells, based on pooled specimen from pre- and post-HIPEC tissues using canonical cell markers shows three distinct populations: monocytes in cluster 4 (*CD14, CD4, FCGR3A, ITGAX, LYZ, VIM, HLA-DRA*) and 18 (*IRF8, and HLA-DRA*); macrophages in cluster 9 (*HLA-DRA, CD14, S100A8, S100A9*); and dendritic cells (DCs) in cluster 11 (*CD4, HLDA-DRA, FCER1A, CLEC10A*) (Figure 2A-2D) (Supplemental Table 2).

Tumor Infiltrating Macrophages (TIM) can polarize and differentiate into two cell types: the anti-tumor classical active macrophages (M1 types) that migrate to tumor sites and suppress oncogenesis (29); and tumor promoter macrophages (M2 type). Our study identified a sub subset of macrophages in cluster 4 (Monocyt_1) that were enriched with *CD163*+ and *CD68*+, markers for M2 (30) (Supplemental Table 2 and Table 6). Moreover, post-HIPEC tissues exhibit a decrease in the density of the M2 population (t test >0.05) (Figure 3B). These observations are significant as M2 macrophages have the ability to modulate extracellular matrix and produce inflammatory cytokines to support tissue regeneration, cancer progression, and metastasis (31). DGE analysis showed a significant decrease in Activating Transcription Factor 3 (*ATF3*) in post-HIPEC tissues (Supplemental Table 5). ATF3 is a member of ATF/CREB family that act to inhibit expression of pro-inflammatory cytokines and control the polarization of macrophages from M1 to M2 phenotypes (32-34).

GSEA analysis indicates that metabolic pathways, including fatty acid and glycolytic pathways, increased in response to HIPEC. These are known to be essential for macrophage activation (Supplemental Figure 3B) (35). We identified Mammalian Target of Rapamycin complex 1 (MTORC1) signaling pathway as significantly enriched in the myeloid cells, which is indicative of increased cellular differentiation of monocytes (36).

### CD74-Enriched Epithelial Ovarian Cancer Cells Demonstrate Activation of Metabolic and Complement Pathways

Epithelial cells were localized in cluster 7 (Figure 2A-D, Supplemental Table 2) and showed no significant changes in cell density pre- and post-HIPEC (Figure 3B). At the transcriptional level there were significant increases in heat-shock proteins, *HSPA6, DNAJB1, CRYAB*, and *HSP90AB1* (21, 22, 37), in the HIPEC-treated specimen (Supplemental Table 7). *HPA6*, in particular, has been previously found to be upregulated in ovarian cancer cell lines treated with Magnetic Fluid Hyperthermia MFH, and its inhibition *in vivo* was successful in reducing tumor growth (38).

DGE analysis indicated a more complex landscape with increased expression of *CD74, CD44*, and *MIF* transcripts (fold change >10) (Supplemental Table 7). The expression of these transcripts in ovarian cancers have already been confirmed at the RNA and protein level (39-42), however, their role in EOC has yet to be investigated. GSEA pathway analysis demonstrated that complement protein and metabolic pathways (fatty acid and glycolysis) are increased in epithelial cells (Supplemental Figure 3C). Interestingly, in ovarian cancer, complement has been found to have non-canonical activity as a mediator between the cancer cells and the tumor microenvironment (TME), including the recruitment of tumor infiltrating immune cells (43, 44)

### Hypoxia and Protein Secretion Signatures in Cancer Associated Fibroblasts (CAFs)

Stromal cells were identified in cluster 8 (*DCN+, PDGFRB+*) and cluster 2 (*DCN+, PDGFRB-*), while cluster 10 represented ovarian myofibroblasts (*DCN+, COL1A1+, COL3A1+*) (Figure 2A2D, Supplemental Table 2). The transcriptomic analysis of these clusters identified significant increased expression of *FAP*, fibroblast activation protein, a marker that specifically labels cancer associated fibroblast (CAF) (45), (p<0.01, fold change >3) (Supplemental Table 8). CAFs represent a cellular subtype that constitutes the stroma in ovarian tumor tissues. The CAF populations show a significant (P < 0.001) decrease in the level of *TCF4* (T-cell transcription factor) that is associated with patient survival in EOC (46). *PTX3* is significantly increased in CAF in post-HIPEC tumors. Several studies report that PTX3 controls tissue regeneration, inflammatory response, and complement activation (11). Differential expression of PTX3 was detected in bulk analysis and here we identify at the single cell resolution that *PTX3* is specifically expressed within the CAF.

Functional GSEA pathway analysis of combined clusters 2, 8, and 10 shows significant increase in activation of protein secretion and hypoxia (Supplemental Figure 3D). Therefore, we screened the DGE list and found a significant elevation in several secretome transcripts that are known to be released by CAF in maintenance of tumor development and response to chemotherapy (47), including *IL-6, I CXCL1, CXCL8, CXCL12*, and *TGFB1* (Supplemental Table 8).

## Discussion

HIPEC treatment leads to a significant and substantial improvement in OS in patients with EOC who undergo optimal interval debulking surgery. However, the manner in which HIPEC exerts this effect remains unknown and poses a barrier to rational use and design of adjunct therapies to potentiate the beneficial effects of HIPEC in these patients. Previous studies investigating HIPEC mechanisms focused on cellular and animal-based models and have not addressed the impact of the tumor microenvironment. We directly address this limitation by applying single cell RNA sequencing technology in matched pre- and post-treatment tumor tissue. Our findings define the landscape of the EOC tumor microenvironment in omental tissue and identify the key molecular and cellular changes that occur directly following the 90-minute treatment period.

We applied an novel strategy to obtain specimen at time of debulking and immediately post-HIPEC treatment. This specimen collection strategy coupled with single cell RNA sequencing allows us to assess the early transcriptomic events of this treatment paradigm. We were able to observe transcriptomic changes within 90 minutes and identify specific cell types where these changes occur. Previous transcriptomic analysis focused on cellular models of cancer and the impact of hyperthermia (9). These studies serve to provide critical insights on the pathways activated by hyperthermia including heat shock proteins (21, 37). Indeed, we observed activation of heat shock response pathways at the bulk RNAseq level. Moreover, we identified activation of EMT and inflammatory pathways within the 90-minute treatment window. Increasing evidence reveals a pivotal role of EMT in tumorigenesis and chemo-resistance (48, 49). With the power of single cell sequencing, we identified the specific cell populations that exhibit an increase in heat shock, EMT, inflammatory pathways, and specific DEGs. We were able to localize *PTX3* upregulation to the CAF population. The function of *PTX3* has not been elucidated in ovarian CAFs, however in breast cancer, inactivation of CEBPD/PTX3 using RI37, a PTX inhibitor peptide, provides benefit in preventing infiltration of drug-resistant cells and limits cancer proliferation (12). Our studies take a leap forward by identifying the transcriptomic changes on a more global scale by providing the transcriptomic changes in specific cell populations impacted by HIPEC within a narrow time window.

Our data reveal that HIPEC triggers activation of adaptive immune and inflammatory responses in T lymphocytes and tumor-infiltrating B lymphocytes (TIL-B). Expansion of activated T-cell populations is achieved by stressing the cells to rapidly replicate and synthesize essential RNA and proteins in a very short time period that can reach 6 hours in some cases (50, 51). Here, we observed these transcriptomic changes are activated within 90 minutes, albeit a short period of time. Further, TIL-Bs in ovarian cancer have recently attracted attention and their presence correlates with improved patient survival (52-56). Our studies complement these findings by demonstrating, at single cell resolution, the existence of a large population of B cells that overexpress the canonical cell markers *MS4A1, BANK1*, and *CD79A*. Moreover, we found transcriptomic activation of estrogen signaling in B cells that determines the fate of cellular activation and production of immunoglobulins in some cancers and autoimmune diseases (57, 58).

Our single cell RNA sequencing analysis revealed that HIPEC leads to reduced M2 macrophages (cluster 4). This is a particularly important observation, especially given the rapid window of our studies, as increasing evidence support the notion that M2 is associated with reduced patient survival, increased chemo-resistance, and poor prognosis (59, 60). In future studies, the M2 macrophages will be investigated directly using flow cytometry analysis to identify these cells and their potential for therapeutic targeting.

We demonstrated that *CD74, CD44*, and *MIF* are transcriptionally upregulated in HIPEC-treated epithelial cells. This is of particular relevance as higher MIF levels have been detected in the sera of EOC patients compared to healthy controls and is was positively associated with overall survival (61). The binding of MIF to its receptor, CD74, stimulates the cleavage of CD74 intracellular domain and modulates signaling pathways to regulate immune response and maintain cellular proliferation and tumorigenesis; this increases with the interaction of CD74 and CD44 (39, 62).

Collectively our data represent the first comprehensive analysis of EOC during HIPEC treatment. We interrogated the cellular and molecular landscape at single cell resolution and reveal insights on the rapid changes that accompany HIPEC. Our findings define the heterogeneity of EOC at the cellular and molecular level. We identified cellular clusters that are enriched in both tumor suppressor and tumor enhancer genes. This indicates different response mechanisms and opportunities to target these pathways and improve patient outcomes. Overall, by mapping the cellular and molecular landscape of HIPEC treated EOC, we provide insights into the rapid changes that accompany HIPEC and provide an opportunity for development of diagnostic and therapeutic targets to augment the patient response to HIPEC and improve patient survival.

## Methods

### Ethical approval and specimen collection

This study was approved by the Institutional Review Board (IRB) of Cleveland Clinic. Five female patients with high grade serous carcinoma were recruited and consented (median age of 64) (Table 1). Tumor specimen was harvested at the time of interval debulking surgery and immediately after HIPEC administration (90 minutes). Specimens were obtained from the same site (omentum) in order to ensure for similar tumor microenvironment.

### Tumor Dissociation

A total of ten tumor specimens were dissociated into a single cell suspension using a Papain Dissociation System (The Worthington Papain Dissociation System, kit# LK003153) according to the manufacturer’s protocol. A single cell suspension of each sample was then stored frozen at −80 C. Prior to sc-RNA sequencing, dissociated samples were thawed in 37^°^C (water bath), centrifuged, and suspended in 10% FBS containing DMEM media, and then sent to Macrogen and the Lerner Research Institute Genomics Core for single cell RNA sequencing.

### Single cell RNA-sequencing analysis

The single-cell RNA-sequencing libraries were generated utilizing the Chromium Single Cell 3’ V3 Kit (10X genomics) according to manufacture instruction. Following library preparation, sequencing was performed using Illumina (NovaSeq6000), and the raw data generated was aligned to GRCh38-3.0.0 using CellRanger 3.1.0. Sequencing depth and cellranger outputs were recorded and included (supplemental table 1). Post data alignment Seurat Version 3.1.5. in R version 4.0 was used for all downstream analysis and data integration. For each cell, the percentage of mitochondrial and ribosomal genes were calculated and then the SCTransform function was utilized to normalize the data and regress out the percent mitochondrial and percent ribosomal genes. Next, cells with more than 6,000 features of RNA and less then 300 features of RNA were removed along with those cells which had greater than 20 percent mitochondrial gene content. After normalization, samples were integrated together using Seurat SCTransform normalization and rpca reduction. Using SCT counts a UMAP was then generated for all samples with overlapping identities. Clusters were subsequently manually identified using top markers of previously published datasets on single cell ovarian cancer (9) additionally top markers of cell populations were used along (10) with expert immunologist verification of population identify using known gene expression of myeloid, T-cell, and epithelial cell markers and their expression or lack of expression on clusters (Supplemental figure 2). When more than one population was present with no obvious or known identifiers, which would constitute this population as a subset, these populations were generically termed with an underscore and a unique number. The abundance of each population was recorded as a percentage of total cells in the sample and then compared in the overall pre vs post analysis. These studies used a paired t-test with Wilcoxon identifying no significant differences in the abundance of cell populations.

### Gene and Pathway Enrichment Analysis

Differential gene expression (DGE) was performed using Seurat comparing all pre-treatment (n=5) vs all post-treatment (n=5) samples with the raw RNA counts with a log fold change threshold of 1. For sub-analyses, the data was subset based on cell identity of interest and then compared pre-treatment vs post-treatment for all samples. Output from this differential expression analysis was then used for Volcano plots using (EnhancedVolcano version 3.11 in R version 4.0). The data output from differential gene expression were then used for Gene Set Enrichment Analysis (GSEA) with the c7.al.v7.1 Hallmark geneset. GSEA was performed by clusterProfiler version 3.11 with the differential gene expression list ordered by adjusted P values and Average Fold Change prior to input and a cutoff p-value of 1. NES and −log10 p-value were then graphed to represent the pathways that are increased post-treatment (Red) and those pathways that were higher pre-treatment (Blue).

### Data Availability

Raw fastq files will be deposited in Gene Expression Omnibus (GEO) Dataset along with the individual cell count matrix of the dataset. In addition, the Seurat Object post data integration including SCT counts will be deposited with cell identities as they are described in this publication.

## Acknowledgments

The authors would like to thank all the members in the Reizes’ laboratory for their insights and contributions during the data acquisition and writing of the manuscript. We would also like to thank members of the flow cytometry and genomic facilities including Drs. Asosingh and Horton. The research was funded by VeloSano Bike to Cure (RD, OR), The Laura J. Fogarty Endowed Chair for Uterine Cancer Research (OR), and Cleveland Clinic Funds.

## Author Contributions

M. P. H. Experimental design, processing tumors, and writing the manuscript

Z. A. Data analysis, organization of results, and writing the manuscript

T. A. Single cell RNA sequencing Data analysis and interpretation, editing the manuscript

C.J.H. Data analysis and interpretation

E.E. Processing tumors and editing the manuscript

T.H. Data analysis and interpretation

J. L. Study design and editing the manuscript

C.M.M. Experimental design and tumor specimen collection

R.D. Conception, tumor specimen collection, administrative support, and writing/editing the manuscript

O.R. Conception and experimental design, administrative support, writing/editing manuscript, and Principal Investigator

## Competing Interests

### Conflicts of Interest

The authors have declared that no conflict of interest exists

## Supplemental Figures

**Supplemental Figure 1.**
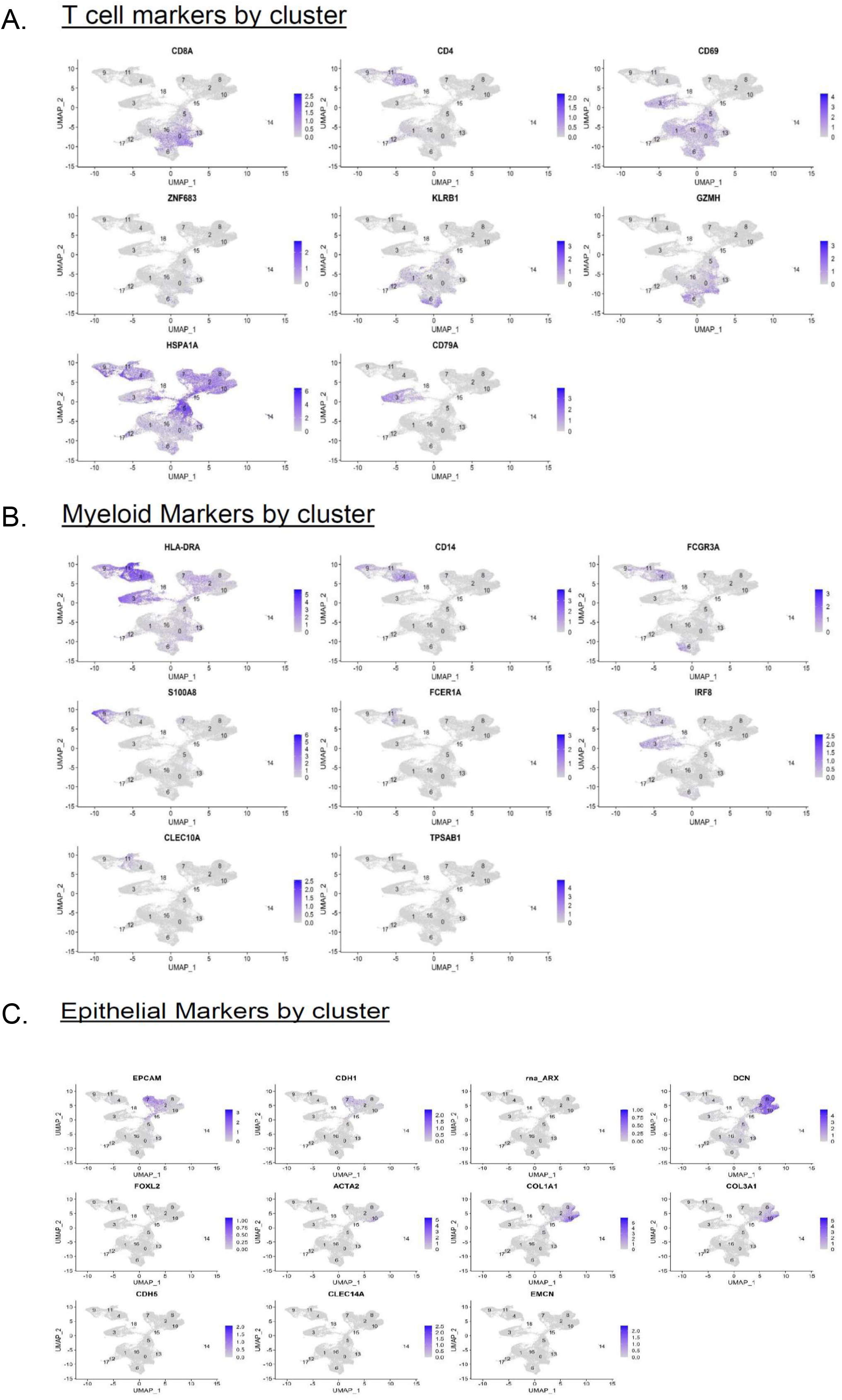
Cellular annotation represented by UMAP plot showing the expression of canonical cell markers in lymphocytes (A) Myeloid cells (B) and Epithelial cells (C)

**Supplemental Figure 2.**
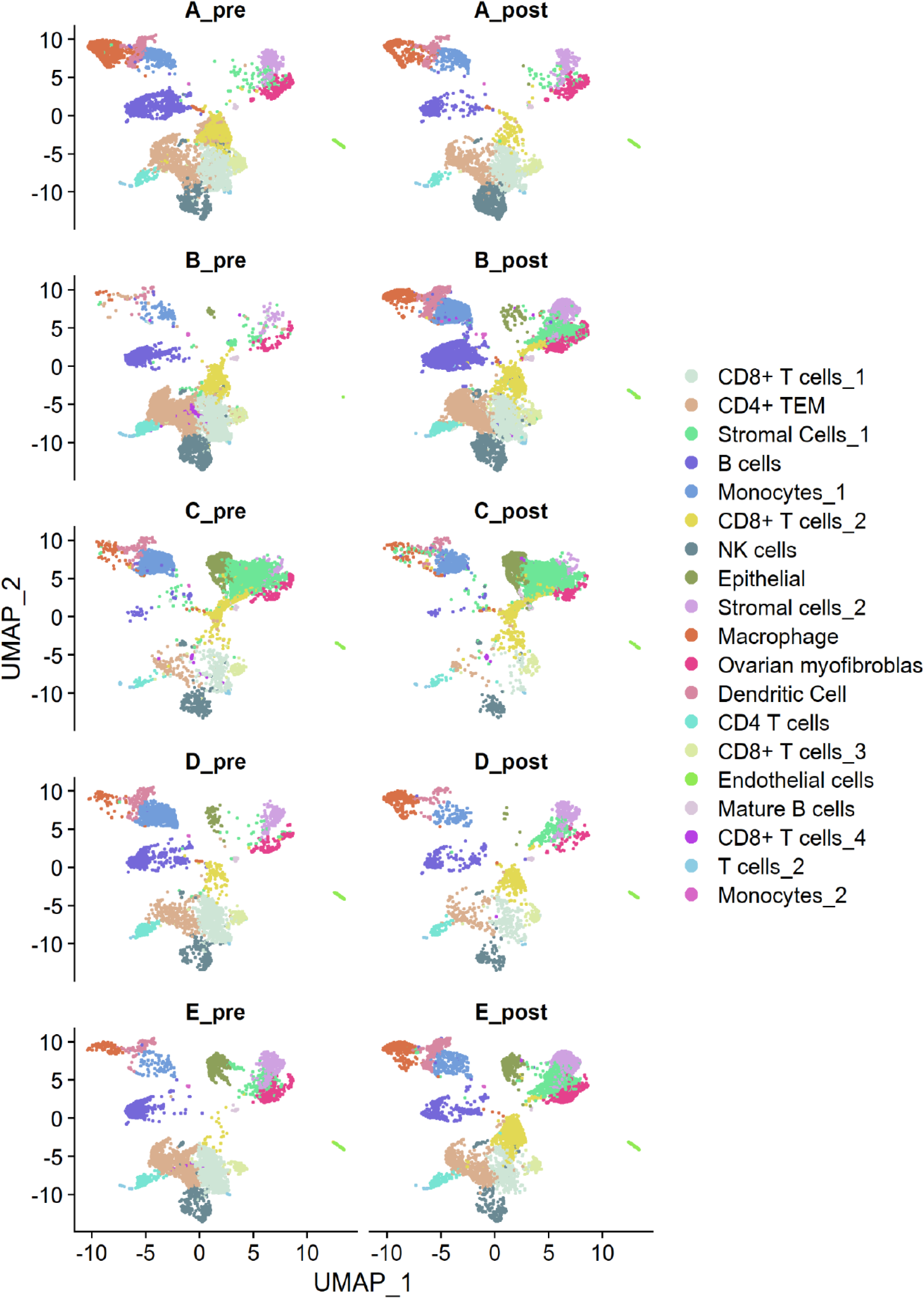
UMAP projections of pre HIPEC sample and matched post HIPEC tissue per patient. The distribution of the identified 19 clusters are represented in UMAP for each individual sample, for patient A, B, C, D and E.Each color represent a distinct cell type distributed in a unique cluster.

**Supplemental Figure 3.**
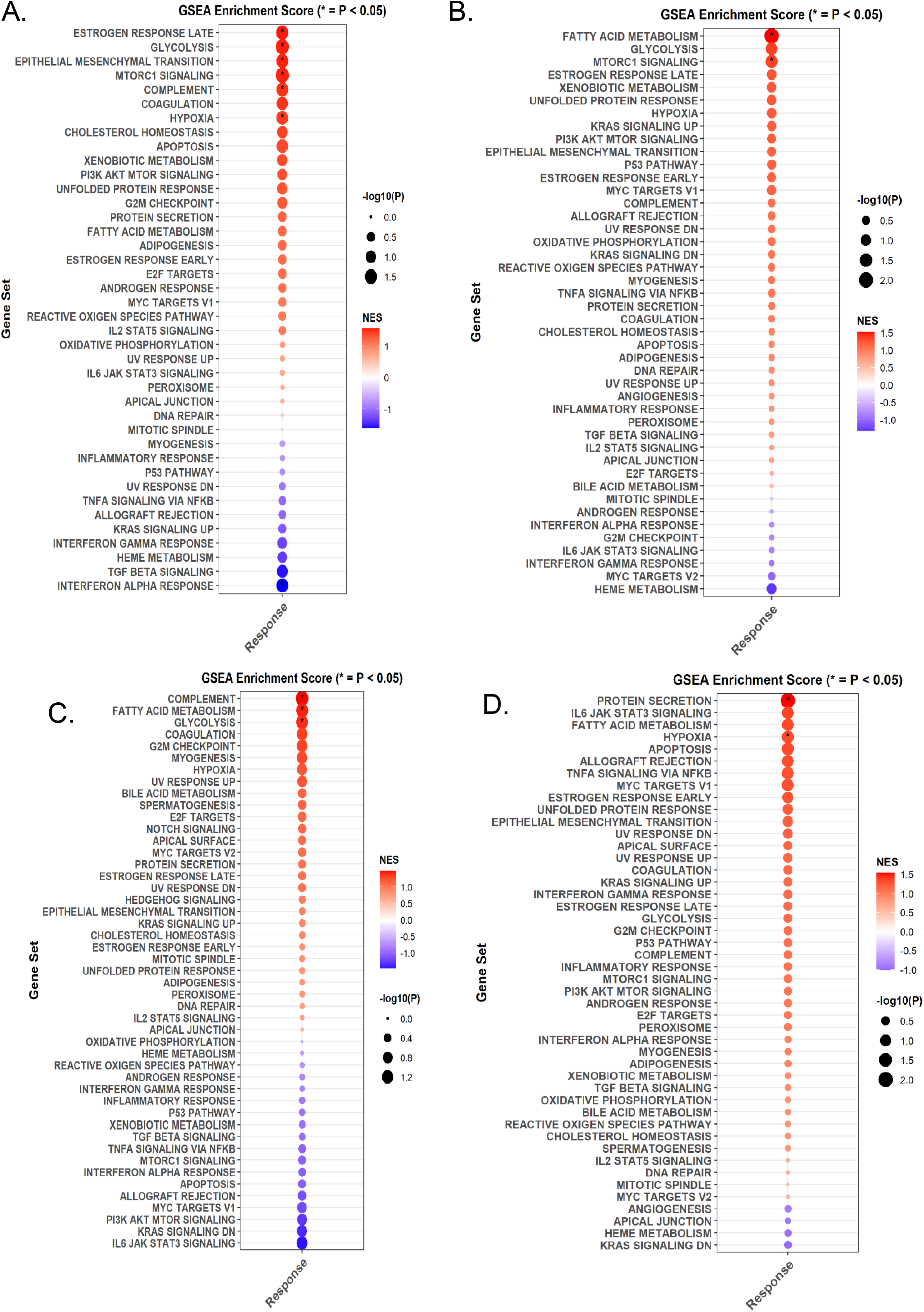
Dotplot representing the functional pathway analysis in post HIPEC tissues vs pre HIPEC samples based on the gene set. (A) B-lymphocytes show upregulation in estrogen response, glycolysis, EMT, and complement pathways. (B) Meyloid cells demonstrated positive stimulation of fatty acid metabolism and mTOR signaling pathway. (C) Epithelial cell population exhibits activation in metabolic and complement pathways. (D) Stromal cells show positive regulation of hypoxia and protein secretion.

## Notes

### Competing Interest Statement

The authors have declared no competing interest.

